# Systematic loss of function screens identify pathway specific functional circular RNAs

**DOI:** 10.1101/2022.10.22.513321

**Authors:** Ling Liu, Matthew Neve, Laura Perlaza-Jimenez, Azelle Hawdon, Simon J. Conn, Jennifer Zenker, Pablo Tamayo, Gregory J. Goodall, Joseph Rosenbluh

## Abstract

Circular RNAs (circRNAs) are covalently closed single stranded RNAs that are produced by RNA back-splicing. A small number of circRNAs have been implicated as functional, however, we still lack systematic understanding of cellular processes and signalling pathways that are regulated by circRNAs. A major gap in understanding circRNA function is the ability to define pathways that are regulated by circRNAs. Here, we generated a pooled shRNA library targeting the back-splice junction of 3,354 human circRNAs that are expressed at low to high levels in humans. We used this library for loss of function proliferation screens in a panel of 18 cancer cell lines from four tissue types that harbour mutations leading to constitutive activity of defined pathways. Using this dataset, we identify context specific and non-specific circRNAs. We validated these observations with a secondary screen and uncovered a role for *circRERE*, a cell essential circRNA that regulates ferroptosis. Furthermore, we characterised the functional roles of pathway-specific circRNA, *circSMAD2*, a novel regulator of the WNT pathway and *circMTO1*, a regulator of MAPK signalling in a *PTEN* dependent manner. Our work sheds light on molecular pathways regulated by circRNAs and provides a catalogue of circRNAs with a measurable function in human cells.

## Introduction

Circular RNAs (circRNAs) were initially described in 1976 by Sanger et al. which visualised by electron microscopy circular RNA molecules that were termed plant viroids^1^. More recently, high throughput RNA-Sequencing (RNA-Seq) in metazoans revealed widespread expression of circRNAs in different organisms and tissue types^2-6^. Initially thought to be an erroneous by-product of splicing, recent evidence suggests that circRNAs are produced by a coordinated and evolutionarily conserved cellular machinery and may have a function in regulating biological processes^5, 7^. For example, during epithelial to mesenchymal transition in mammary cells, QKI, an RNA splicing regulator, mediates circRNA formation in a coordinated manner^8^. Further evidence that circRNAs play a role in regulating biological processes include large scale circRNA expression profiling across healthy and cancer tissues^2^. These studies found that circRNAs expression is not correlated with linear mRNA expression suggesting that circRNAs have distinct functions. Moreover, functional studies also provide evidence that some circRNAs have a function. Short hairpin RNA (shRNA) based screens limited to highly expressed circRNA in prostate cancer cell lines found a subset of prostate cancer dependent circRNAs^9^ and a mouse knockout of circular *cdr1as* showed a neurological phenotype^10^. Taken together, these observations suggest that at least some circRNAs are functional and have a role in regulating biological processes. However, no large-scale systematic investigation of circRNAs in the context of pathway specific activity has been done.

The WNT and MAPK signalling pathways are key regulators of development, growth and differentiation and are deregulated in various cancer types^11-13^. Understanding the cellular components that regulate the WNT and MAPK pathways in normal and disease conditions, has enhanced our understanding of fundamental biological processes and has enabled new strategies for patient therapy. Specifically, studies in cultured cell lines and animal models revealed proteins that directly or indirectly regulate these pathways. Based on these studies, various drugs have been developed. These include, MEK1 inhibitors^14^ that are used in cancers with deregulated MAPK signalling and Porcupine inhibitors that are currently in clinical trials for cancers with deregulation of WNT signalling^15, 16^. However, in contrast to the detailed studies of proteins that regulate these pathways, we lack comprehensive understanding of how non-coding elements such as circRNAs may regulate these key signalling networks.

Loss of function phenotypic screens in cancer cell lines have been highly valuable for defining genes that are associated with signalling pathways^17, 18^. Using genomic profiles (e.g. mutations, copy number alterations and gene expression changes) in hundreds of cell lines enable classification of activated signalling pathways^19, 20^. Since cancer cell lines require constitutive activity of these signalling pathways, identifying genes that are required for proliferation only in the context of an activated pathway could identify regulators of these pathways. This strategy has been successfully used for identifying genes that regulate signalling networks^17, 18, 21^.

Here, we use a similar strategy to identify circRNAs that are important for the activity of the WNT and MAPK signalling pathways as well as common essential and tissue specific circRNAs. We show that although most circRNAs do not have a functional phenotype in this assay system, some circRNAs are direct regulators of important signalling pathways.

## Results

### Selection of circRNAs for loss of function screens

Using exome capture RNASeq Vo et al. profiled circRNA expression across human cancer and normal tissues and cell lines in >2,000 samples^2^. Since circRNAs are highly stable^5^, it is possible that, even when expressed at very low levels they are still functional. Therefore, for functional screens we selected every circRNA that was detected by Vo et al.^2^ with at least 30 reads in at least 10 samples. This resulted in 3,354 circRNAs that were used in all subsequent screens (Fig. 1A and Supplementary Table 1).

**Figure 1:**
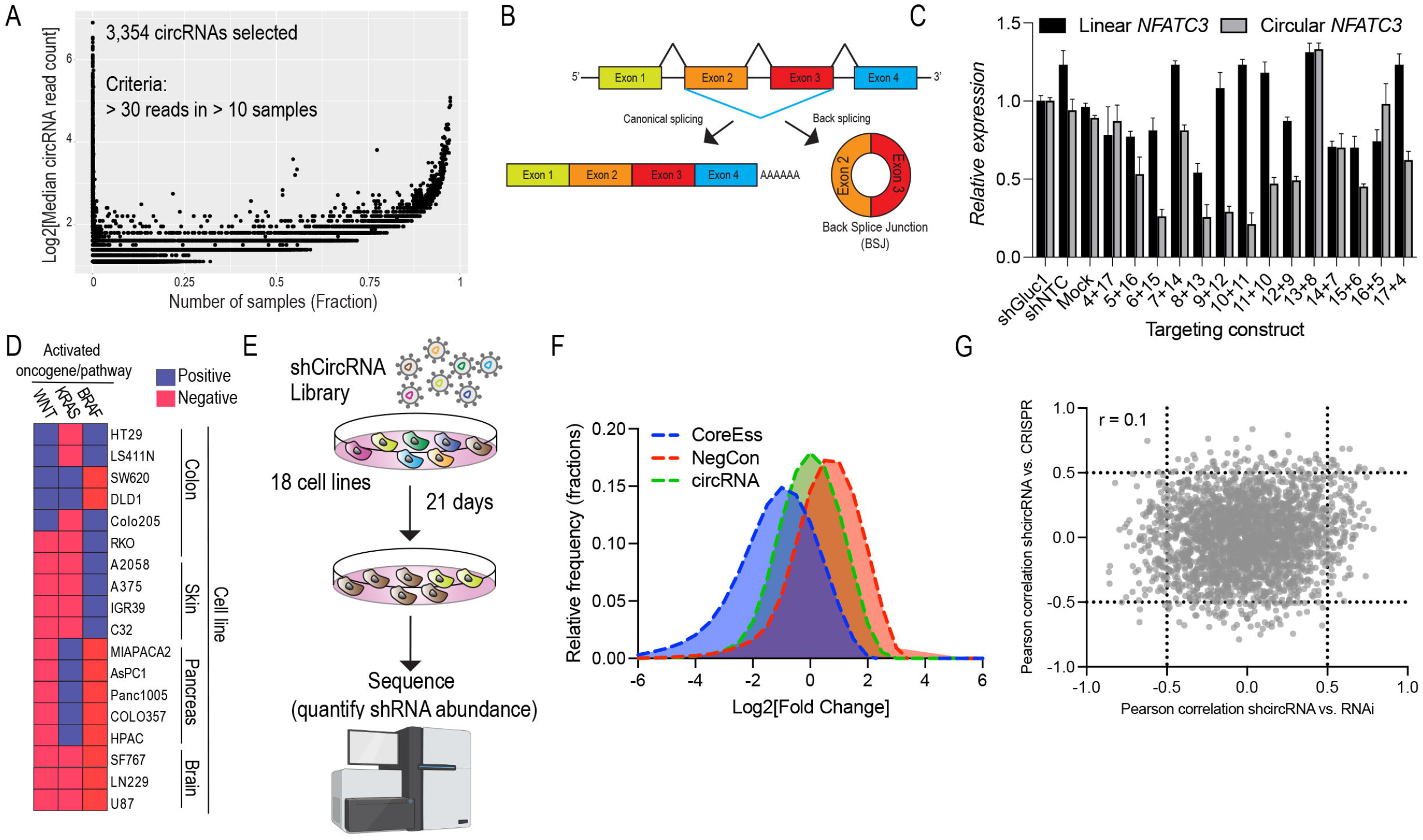
Primary loss of function screens identify circRNA dependencies. (A) Expression of all circRNAs identified by Vo et al.^2^. For functional screens we selected circRNAs that were found in ≥10 samples with ≥30 reads. (B) Depiction of the circRNA back-splice junction (BSJ) that is used for design of circRNA targeting shRNAs. (C) Optimisation of shRNAs. HEK293T cells were transfected with shRNAs targeting the BSJ of *circNFATC3*. Numbers represent nucleotides from each side of the BSJ. 48h post transfection qRT-PCR was used to quantify *circNFATC3* and linear *NFATC3* RNA. (D) Cell lines used in primary screen and pathways dependencies in these cell lines. (E) Description of primary loss of function screen. (F) Distribution of all shRNAs targeting, positive controls (core essential genes) negative controls (non-targeting shRNAs) and circRNAs. (G) For each circRNA available data from DepMap was used to calculate correlation between phenotype of circRNA knockdown and CRISPR or shRNA suppression of its gene precursor.

Previous studies have shown that shRNAs targeting the circRNA back-splice junction (BSJ) result in circRNA suppression without having any effect on the linear mRNA transcript (Fig. 1B and^9^). To assess the extent of overlap that is necessary for effective circRNA suppression, we used qRT-PCR with primers unique to *circNFATC3* or linear *NFATC3* to measure circular or linear *NFATC3* levels following introduction of shRNAs targeting the *circNFATC3* BSJ in HEK293T cells (Fig. 1C). We found that 21-mers with an overlap of at least 8nt on each side of the BSJ result in efficient circRNA knockdown with little effect on the mRNA. Based on these observations we designed a circRNA-targeting shRNA pooled library. For each circRNA we designed 5 shRNAs and we also included 597 shRNAs targeting 201 core essential genes^22^ and 722 negative control shRNAs (Supplementary Table 2). Compared to previously described circRNA loss of function screens^9^ our library contained 939 circRNAs that were used in previous screens, but most of our library (2,415, 71%) contained shRNAs targeting circRNAs that have not been screened before (Supplementary Fig. 1A).

### circRNA loss of function screens

For functional screens we selected 18 cancer cell lines of 4 tissue types (colon, pancreas, skin and brain) that have different mutation profiles that result in pathway specific dependencies (Fig. 1D and^17, 22^). For example, cell lines with mutations in the tumour suppressor *APC* or the oncogene *CTNNB1* have constitutive activation of the WNT signalling pathway and require the expression of WNT pathway components for proliferation^20^. Similarly, proliferation of cell lines with activating *BRAF* mutations are dependent on components of the RAF/MAPK pathway^23^. Cells were infected with the pooled circRNA targeting shRNA library at 1,000 cells/shRNA and low multiplicity of infection (MOI = 0.3) to ensure one shRNA/cell. After 21 days in culture DNA extracted from these cells was used for sequencing and quantification of shRNA abundance (Fig. 1E and Supplementary Table 3). As expected, shRNAs targeting known core essential genes^22, 24, 25^ had a negative impact on proliferation and negative controls centred around zero (Fig. 1F and Supplementary Fig. 1B). To identify circRNAs that are required for proliferation of a particular cell line we calculated the average Log2[Fold change] of the five shRNAs used to target a circRNA to generate a circRNA score (Supplementary Table 4). We found, that although most circRNA knockdowns did not have any effect on proliferation some circRNA knockdowns had an impact on cell proliferation (Fig. 1F), suggesting these are functional circRNAs that regulate biological processes in a context specific or non-specific manner.

Previous studies showed that the expression of most circRNAs is not correlated to the expression of their linear precursor^2, 9^. To evaluate the correlation between linear and circRNA dependency, we obtained from the Depmap portal CRISPR^26^ or RNAi^27^ gene dependency scores for the linear precursor of each of the circRNAs we tested. By comparing gene and circRNA dependencies, we found that circRNA dependencies are not correlated with gene dependencies (r = 0.1)(Fig. 1G and Supplementary Table 5).

Using the circRNA score we defined different types of functional circRNA (Supplementary Table 6). As core essential circRNAs we classified circRNAs that were essential (Log2[Fold change] < -1) in ≥ 94% (17 cell lines) of the cell lines we screened. This resulted in 89 core-essential circRNAs (2.5% of all screened circRNAs). To identify circRNAs that function in a context/pathway specific manner, we used a two-class comparison to compare circRNA dependencies in cell lines that are dependent on known signalling pathways or lineage dependencies (Supplementary Table 6). Specifically, we classified cells as either WNT, *KRAS* or *BRAF* active and identified circRNAs that are required for proliferation in cell lines that harbour activating mutation in these pathways (Supplementary Table 6). Similarly, we identified circRNAs that are tissue specific (brain, pancreas, skin, colon) (Supplementary Table 6). Since most cancer specific dependencies are enriched in particular linages (e.g. WNT in colon and KRAS in pancreas) it is difficult to delineate between a pathway specific or tissue specific dependency. Thus, we conducted secondary screens in order to validate these observations.

### Secondary validation screen

To validate these observations of core essential and pathway specific circRNAs, we conducted a secondary validation screen in a panel of 11 cancer cell lines. From each two-class comparison analysis we selected the top 100 scoring circRNAs (based on the delta score or the number of dependent cells for core essential circRNAs) resulting in a total of 480 circRNAs (Fig. 2A). For each circRNA we designed 5 shRNAs. In addition, we included as positive controls shRNAs targeting core essential genes and shRNAs targeting known components of signalling pathways (Supplementary Table 7). Following transduction cells were propagated for 21 days and genomic DNA was used for quantification of shRNA abundance (Supplementary Table 8). Using the average Log2[fold change] (Supplementary Table 9) we calculated a circRNA score in each cell line. As expected, depletion of core essential genes had a dramatic effect on cell proliferation (Supplementary Fig. 2). Since Log2[Fold change] values for core essential controls were centred around -1 for most cell lines (Supplementary Fig. 2), as validated core essential circRNAs we considered circRNAs that had a circRNA score ≤ -0.5 in at least 7 cell lines (Fig. 2B). Using this approach 20 circRNAs that scored in both primary and secondary screens were implicated as strong core essential circRNAs.

**Figure 2:**
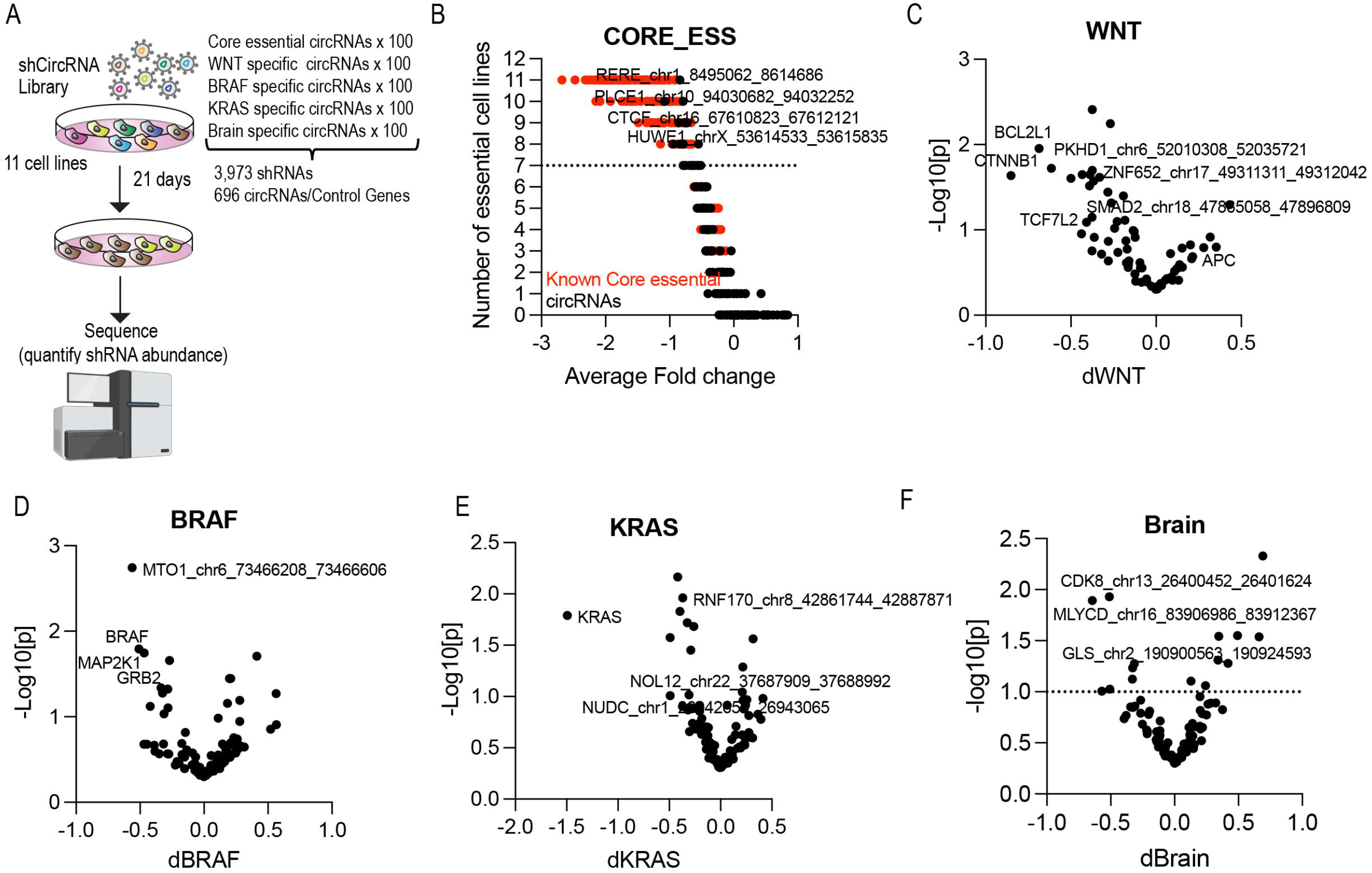
Secondary validation screen defines context dependent and independent circRNAs. (A) Secondary screen schematics. Top 100 circRNAs from each two-class comparison were selected for secondary screens. (B) Validation of core essential circRNAs. (C) Validation of WNT specific circRNAs. (D) Validation of *BRAF* specific circRNAs. (E) Validation of *KRAS* specific circRNAs. (F) Validation of brain specific circRNAs.

To identify pathway specific circRNA dependencies we compared the circRNA dependency scores between cells with or without constitutive activity of WNT (Fig. 2C), *BRAF* (Fig. 2D) or *KRAS* (Fig. 2E). Since circRNAs have been shown to be highly active in brain cells^28^, we also included brain specific circRNAs and brain cancer cell lines (Fig. 2F, and Supplementary Table 10). As expected, known components of these signalling pathways scored as pathway specific dependencies. Specifically, WNT pathway genes such as, *CTNNB1, TCF7L2* and *BCL2L1*^*17, 20*^ scored in WNT driven cells (Fig. 2C). Components of the MAPK signalling pathway *BRAF, MAP2K1* and *GRB2* scored as differentially essential in *BRAF* driven cells (Fig. 2D) and *KRAS* scored as the most differentially required gene in *KRAS* mutant cell lines (Fig. 2E). Taken together, these results provide a list of high confidence core essential as well as pathway specific functional circRNAs.

### Core essential circRNAs

Of the 3,554 circRNAs we assessed in the primary screen the majority (70%) were non-essential (defined as essential in ≤ 2 cell lines). Of the 1,015 remaining circRNAs 926 (28.5%) showed dependency in a defined population of cell lines (between 3-17) and only 89 (2.5%) were essential in ≥ 17 cell lines (Fig. 3A). This most likely reflects the fact that most circRNAs are expressed at low levels and are not functional. Of the circRNAs we identified as core essential, 20 circRNAs validated in our secondary validation screen (Supplementary Table 10). We further validated two of these core essential circRNA dependencies, *circHUWE1 and circRERE*.

**Figure 3:**
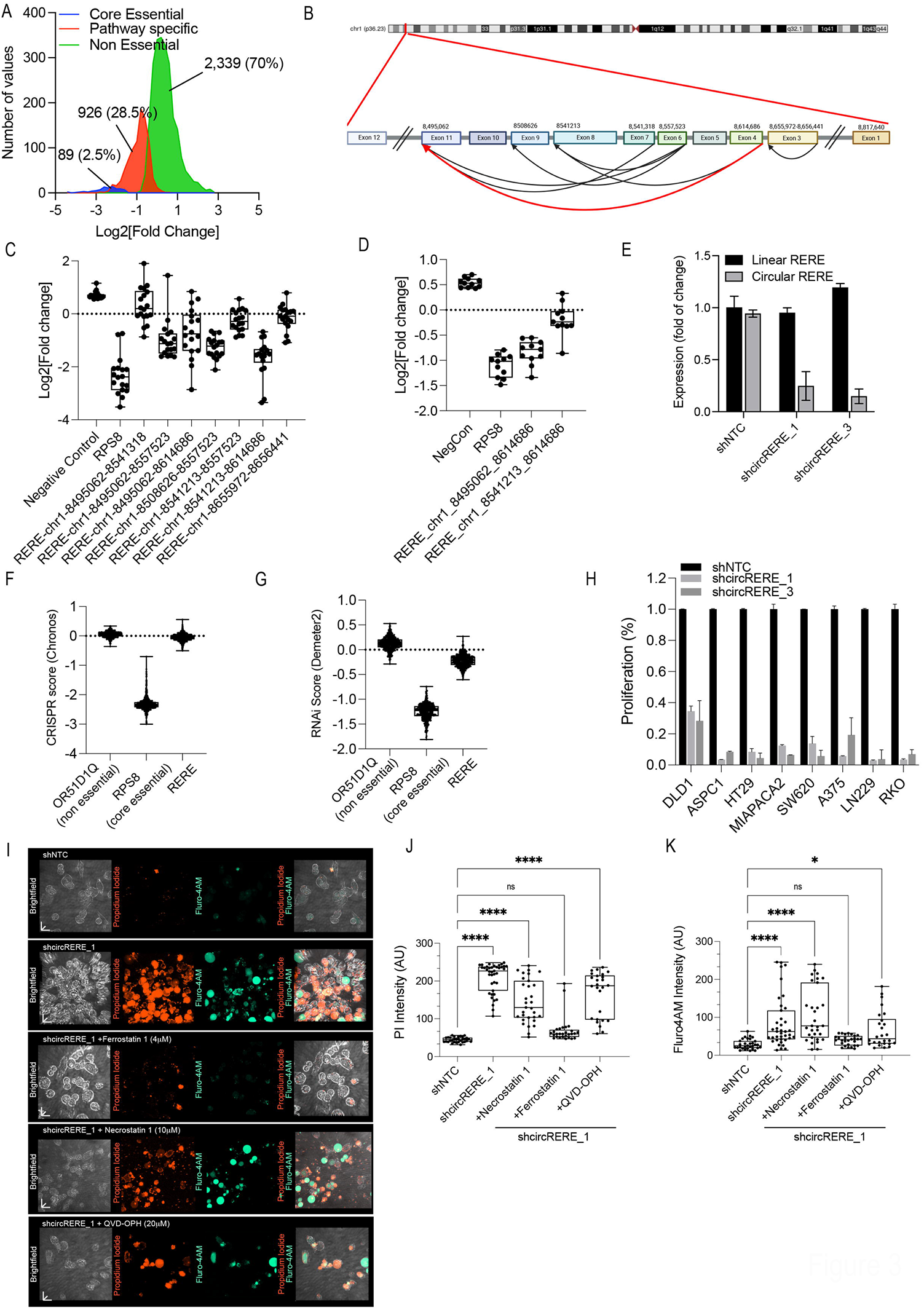
Identification of core essential circRNAs. (A) Distribution of proliferation changes for different types of circRNAs. (B) Genome structure and circRNAs produced from *RERE* gene. (C) Proliferation changes in primary screen induced by shRNAs targeting *circRERE* isoforms. Each point represents a different cell line. (D) Proliferation changes in secondary screen following shRNA mediated suppression of two *circRERE* variants. Only RERE_chr1_8495062_8614686 scored as a cell essential circRNA. Each dot represents a different cell line. (E) Expression of linear or circular circRERE (RERE_chr1_8495062_8614686) three days post transduction with two circRERE targeting shRNAs in DLD1 cells. (F) Proliferation changes from Depmap^51^ following CRISPR mediated suppression of *RERE* gene. (G) Proliferation changes from Depmap^51^ following shRNA mediated suppression of *RERE* gene. (H) Proliferation changes induced by shRNAs targeting *circRERE*. (I) RKO cells were transduced with control or *shcircRERE_1* following selection cells were treated with either 4μM of Ferrostatin-1 or 10μM of Necrostatin or 20 μM of QVD-OPH and cell viability was measured by staining with propidium iodide (PI) or Fluro4AM after 48h. (J) Quantification of PI staining. (K) Quantification of Fluro4AM staining.

Of the two expressed *HUWE1* derived circRNAs (Supplementary Fig. 3A) we found one circRNA (HUWE1-chrX-53614533-53615835) that was essential in all cell lines (Supplementary Fig. 3B). We validated these results in our secondary validation screen (Supplementary Fig. 3C). We further confirmed that the shRNAs we used only supressed *circHUWE1* expression and not linear *HUWE1* (Supplementary Fig. 3D). To further validate these observations in a singleton experiment we used a crystal violet proliferation assay and confirmed that suppression of *circHUWE1* had a negative impact on proliferation in all the cell lines we tested (Supplementary Fig. 3E,F). Interestingly, CRISPR and RNAi experiments show that similar to c*ircHUWE1, HUWE1* gene is a core essential gene that is required for cell survival (Supplementary Fig. 3G,H). This raises the possibility of a mechanism by which *circHUWE1* functions to enhance the activity of HUWE1 protein. This is consistent with the conflicting literature suggesting different functional roles for *circHUWE1*^*29-32*^. HUWE1 is an E3 ubiquitin ligase that regulates different proteins in a context specific manner. Similarly, it is likely that *circHUWE1* regulates essential biological processes in a context specific manner.

The gene *RERE* expresses 7 circRNA isoforms (Fig. 3B) that were included in our primary screen. Two of these *RERE* derived circRNAs were essential across most cell lines that were included in the screen RERE-chr1-8495062-8557523 and RERE-chr1-8541213-8614686 (Fig. 3C). Of these RERE-chr1-8541213-8614686 also validated in our secondary validation screen (Fig. 3D). The magnitude of dependency in these experiments was similar to what we observed following knockdown of *RPS8*, a known core essential gene, indicating a high degree of cell death (Fig. 3C,D). Using qRT-PCR we found that *circRERE* targeting shRNAs only inhibited expression of *circRERE* and had no effect on the expression of *RERE* gene (Fig. 3E). Furthermore, data from Depmap showed that CRISPR (Fig. 3F) or RNAi (Fig. 3G) mediated suppression of *RERE* gene had no effect on cell proliferation indicating that the effect of *circRERE* is unique to one *RERE* circle and is not related to the function of the *RERE* gene. We validate the proliferation effect observed by suppression of *circRERE* using a crystal violet proliferation assay (Fig. 3H and Supplementary Fig. 3I).

To determine the mechanism of cell death following suppression of *circRERE*, we used inhibitors of different cell death pathways to rescue cell death induced by *circRERE*-targeting shRNAs. Treatment with a pan-caspase inhibitor (QVD-OPH) or a Necroptosis inhibitor (Necrostatin) had no effect on the ability of *circRERE*-targeting shRNAs to induce cell death (Fig. 3I-K). However, treatment with the ferroptosis inhibitor Ferrostatin-1^33^ rescued cell death induced by *circRERE* targeting shRNAs (Fig. 3I-K). We further validated these results by measuring cell viability with a trypan blue staining assay (Supplementary Fig. 3J). Ferroptosis is a cell death mechanism induced by iron that is inhibited by *GPX4*, which is required for maintenances of a drug resistant state in cancer cells^34^. These results suggest that *circRERE* is a regulator of ferroptosis and are consistent with previous results that suggest *circRERE* plays a role in regulating drug resistance in multiple myeloma^35^.

### circSMAD2 regulates the WNT signalling pathway

We have previously shown the effectiveness of loss of function shRNAs^17^ or CRISPR^20^ screens, for identification of genes associated with the WNT signalling pathway. To identify circRNAs that have a role in regulation of the WNT pathway we compared circRNA dependencies between WNT active and WNT inactive cell lines (Supplementary Fig. 1C, and Supplementary Table 6). Based on the dWNT score, we selected the top 100 circRNAs for secondary screening (Fig. 2C). As validated WNT dependent circRNAs we defined circRNAs with a dWNT score ≤ -0.2 and a pValue ≤ 0.15 in the secondary screen (Supplementary Table 10). Using these criteria, 18 circRNAs scored as WNT specific dependencies (Supplementary Table 10).

To further validate these observations, we cloned shRNAs targeting three of the top WNT specific hits; *circSMAD2, circPKHD1* and *circZNF652*. Following transduction of WNT active or inactive cell lines we used crystal violet proliferation assay to measure the effect of circRNA knockdown. Confirming the results from our secondary screen, knockdown of these circRNAs inhibited proliferation only in WNT active cell lines (Fig. 4A and Supplementary Fig. 4A-E). We found that shRNAs supressing these circRNAs had a significant effect on proliferation only in WNT active cell lines (*circSMAD2* p<0.0001, *circZNF652* p<0.0001, *circPKHD1* p=0.0327, Fig. 4B). We validated that these shRNAs only supressed circRNA expression without affecting the expression of the linear mRNA using qRT-PCR with primers specific for circular or linear expression (Fig. 4C and Supplementary Fig. 4F,G). Of these circRNAs, *circSMAD2* knockdown had the greatest effect on proliferation in WNT active cancer cells and was chosen for further studies.

**Figure 4:**
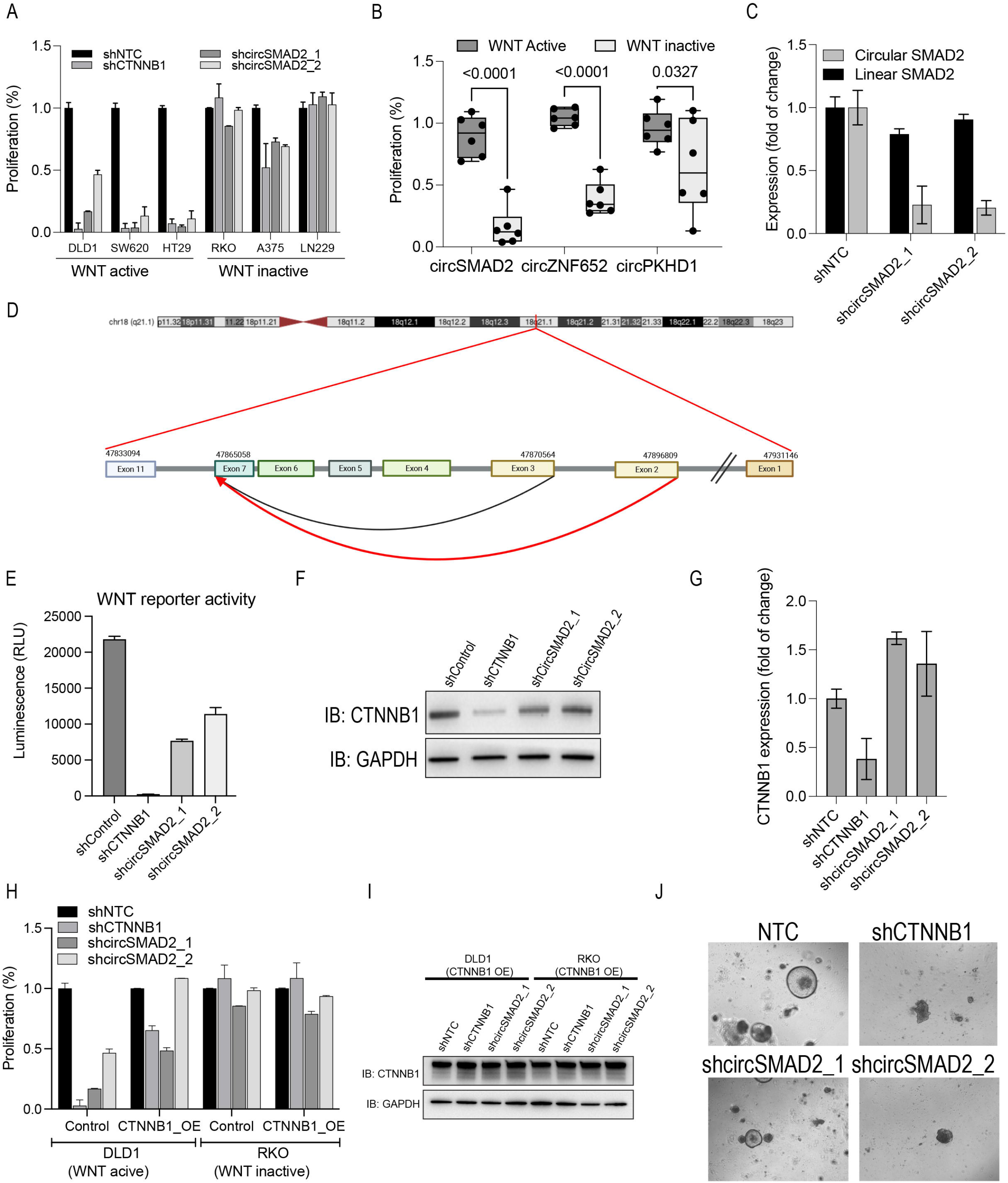
Identification of circRNAs that regulate the WNT signalling pathway. (A) Proliferation of WNT dependent and independent cancer cell lines 7 DPI with shRNAs targeting *circSMAD2*. (B) Proliferation changes in WNT active and WNT inactive cell lines following suppression of *circSMAD2, circZNF652* and *circPKHD1*. p-Value was calculated using one-way Anova. (C) Expression of circular and linear *circSMAD2* in DLD1 cells 3 DPI with circSMAD2 targeting shRNAs. (D) Genome structure and circRNAs produced from the *SMAD2* gene. (E) DLD1 cells stably expressing a WNT reporter were infected with shRNAs targeting *CTNNB1* or *circSMAD2* and WNT activity was measured 5 DPI. (F) CTNNB1 protein levels following expression of shRNAs targeting *circSMAD2*. (G) mRNA levels of *CTNNB1* following expression of *circSMAD2* targeting shRNAs in DLD1 cells. (H) Rescue of circSMAD2 mediated proliferation arrest in cells overexpressing *CTNNB1*. (I) Rescue of *circSMAD2* mediated decrease in CTNNB1 protein levels in cells overexpressing *CTNNB1*. (J) Colon organoid cultures (colonoids) 7 DPI with shRNAs targeting *circSMAD2*.

*SMAD2* mRNA produces two circRNAs (Fig. 4D) that are detected in ≥ 10 samples with ≥ 30 reads (Supplementary Table 1). In samples that express either of the *SMAD2* circRNAs they are found at similar expression levels (average of 64-65 reads). However, of the 2,000 samples profiled by Vo et al.^2^, SMAD2_chr18_47865058_47870564 is expressed in 18 samples and SMAD2-chr18-47865058-47896809 is expressed in 226 samples (Supplementary Table 1). Consistent with its expression pattern in a large number of samples only SMAD2-chr18-47865058-47896809 scores as a WNT dependent circRNA (Supplementary Table 6). The WNT specific proliferation effect we observed following knockdown of *circSMAD2* could be due to a direct or indirect effect on the WNT signalling pathway. To assess the effect of *circSMAD2* knockdown on the WNT signalling pathway we used a luciferase WNT reporter activity assay^36^ in DLD1, a WNT active, cell line. As expected, suppression of *CTNNB1*, the main transcription factor regulated by the WNT pathway^11^, inhibited WNT reporter activity (Fig. 4E). Similarly, we found that suppression of *circSMAD2* expression resulted in a 50% reduction in WNT reporter activity (Fig. 4E). To assess the effect of *circSMAD2* on the WNT signalling pathway we measured *CTNNB1* protein stability following suppression of *circSMAD2*. Consistent with what we observed with the WNT reporter assay, knockdown of *circSMAD2* resulted in a 50% reduction in CTNNB1 protein levels, indicating that *circSMAD2* is directly regulating the activity of the WNT signalling pathway (Fig. 4F). However, suppression of *circSMAD2* did not inhibit *CTNNB1* mRNA levels (Fig. 4G) suggesting, that the effect of *circSMAD2* on the WNT signalling pathway is not due to direct regulation of *CTNNB1* mRNA.

To further validate these observations and to ensure that the phenotypes we observed following *circSMAD2* knockdown are a direct consequence of *circSMAD2* effecting the WNT signalling pathway, we performed a rescue experiment. We overexpressed a mouse *CTNNB1* mutant that lacks serine and threonine residues that are regulated by the destruction complex (S33A, S37A, T41A, S45A)^36^ in a WNT dependent (DLD1) and WNT independent (RKO) cell. Importantly, mouse *CTNNB1* is identical in its amino acids to human CTNNB1 protein but different in its nucleotide sequence and is therefore, not inhibited by shRNAs targeting human *CTNNB1*. Following transduction with *CTNNB1* or *circSMAD2* shRNAs we measured proliferation using a crystal violet staining assay. Overexpression of *CTNNB1* rescued the proliferation inhibition mediated by shRNAs targeting *CTNNB1* and *circSMAD2* (Fig. 4H and Supplementary Fig. 4H) and the reduction in CTNNB1 protein levels (Fig. 4I). These results, suggest that the proliferation inhibition mediated by *circSMAD2* knockdown in WNT dependent cells is due to its effect on the WNT signalling pathway.

As an additional experimental system, we assessed *circSMAD2* as a regulator of WNT signalling in a mouse colon organoid (colonoids) model^37^. Colonoids are made from mouse colonic crypts grown in a Matrigel matrix in the presence of ligands that regulate the WNT, MAPK and TGFβ pathways and require constitutive activity of the WNT signalling pathway. Following infection with shRNAs mediating suppression of *CTNNB1* or *circSMAD2* colonoids were plated in Matrigel and visualized after 7 days. As expected, *CTNNB1* targeting shRNAs abolished the ability of colonoids to grow (Fig. 4J). Similarly, suppression of *circSMAD2* resulted in impaired colonoid growth (Fig. 4J). Taken together, these results demonstrate that *circSMAD2* is a regulator of WNT activity and is essential for survival of WNT driven cells.

### circMTO1 regulates the MAPK signalling pathway

MAPK signalling is regulated by the protooncogenes *BRAF* and *KRAS*^*38*^. *KRAS* mutated cancers have a complex co-dependency profile that involves feedback loops and activation of additional signalling pathways. In contrast, cancers harbouring *BRAF* mutations have constitutive MAPK activity and require MAPK activity for proliferation in cell lines, animal models and humans^39^. To identify circRNAs that are important for regulation of the MAPK signalling pathway, we used a two-class comparison analysis and identified circRNA dependencies that are unique to cell lines with activating *BRAF* or *KRAS* mutations (Supplementary Table 6 and Supplementary Fig. 1D). We selected the top 100 circRNAs for secondary screening (Fig. 2D,E and Supplementary Table 10). As validated circRNA dependencies we considered circRNAs with a dBRAF or dKRAS ≤ -0.2 and a pValue ≤ 0.15 in the secondary screen. Using these criteria, we identified 7 *BRAF* specific circRNAs and 19 *KRAS* specific circRNAs (Supplementary Table 10).

Using a crystal violet staining proliferation assay, we validated *circRNF170* (Supplementary Fig. 5A-C) *circNOL12* (Supplementary Fig. 5D-F) and *circNUDC* (Supplementary Fig. 5G-I) three of the top *KRAS* specific circRNA dependencies in 6 cancer cell lines. Confirming the screen results we found that shRNAs supressing the expression of *circRNF170* or *circNOL12* had a significant effect on proliferation of *KRAS* mutated cancers (p=0.0827 for *circRNF170* and *circNOL12* – Supplementary Fig. 5J), although *circNUDC* shRNAs affected both the *KRAS* mutant and WT cell lines. None of these circRNAs have been described before as functional in any context suggesting *circRNF170* and *circNOL12* as novel KRAS dependencies. To identify possible mechanisms of these *KRAS* specific circRNAs we measured changes in phosphorylation of pAKT and pERK1/2, following shRNA mediated suppression of *circRNF170* and *circNOL12* (Supplementary Fig. 5K,L). We did not observe any changes in these proteins suggesting that these circRNAs regulates alternative *KRAS* pathways.

*circMTO1* (Fig. 5A) scored as the top *BRAF* specific dependency and had a differential effect with a similar magnitude as *BRAF* (Fig. 2D). We validated these results, by cloning these shRNAs and transducing *BRAF* WT or mutant cells. We found that shRNAs targeting *circMTO1* had a dramatic effect on proliferation of *BRAF* mutated cell lines and no effect on *BRAF* WT cells (Fig. 5B,C and Supplementary Fig. 6A). *circMTO1* targeting shRNAs had no effect on the abundance of *MTO1* mRNA and only inhibited the expression of *circMTO1*, indicating the specificity of these *circMTO1* targeting shRNAs (Fig. 5D). To assess if *circMTO1* is a direct or indirect regulator of the MAPK signalling pathway we measured the additive effect of supressing circMTO1 in combination with a BRAF inhibitor on proliferation of *BRAF* mutated cells. *BRAF* mutated cell lines expressing *circMTO1* targeting shRNAs were treated with PLX4032, a potent BRAF inhibitor^40, 41^, and proliferation was measured using a crystal violet proliferation assay 10 days after treatment (Fig. 5E and Supplementary Fig. 6B,C). We found no additive proliferation effect following simultaneous inhibition of BRAF and *circMTO1* indicating that both work in a similar mechanism of action. Furthermore, we found no additive change in pERK1/2 following combined inhibition of BRAF and *circMTO1* (Fig. 5F) suggesting that *circMTO1* is a direct regulator of the BRAF/MAPK signalling pathway. To confirm the effect of *circMTO1* on MAPK signalling we measured the effect of *circMTO1* shRNAs on the known downstream MAPK target, CCND1, and found that while suppression of *circMTO1* did not affect the levels of *BRAF* mRNA, it inhibited expression of *CCND1* (Fig. 5G). Taken together these results suggest that *circMTO1* is a regulator of the BRAF/MAPK signalling pathway.

**Figure 5:**
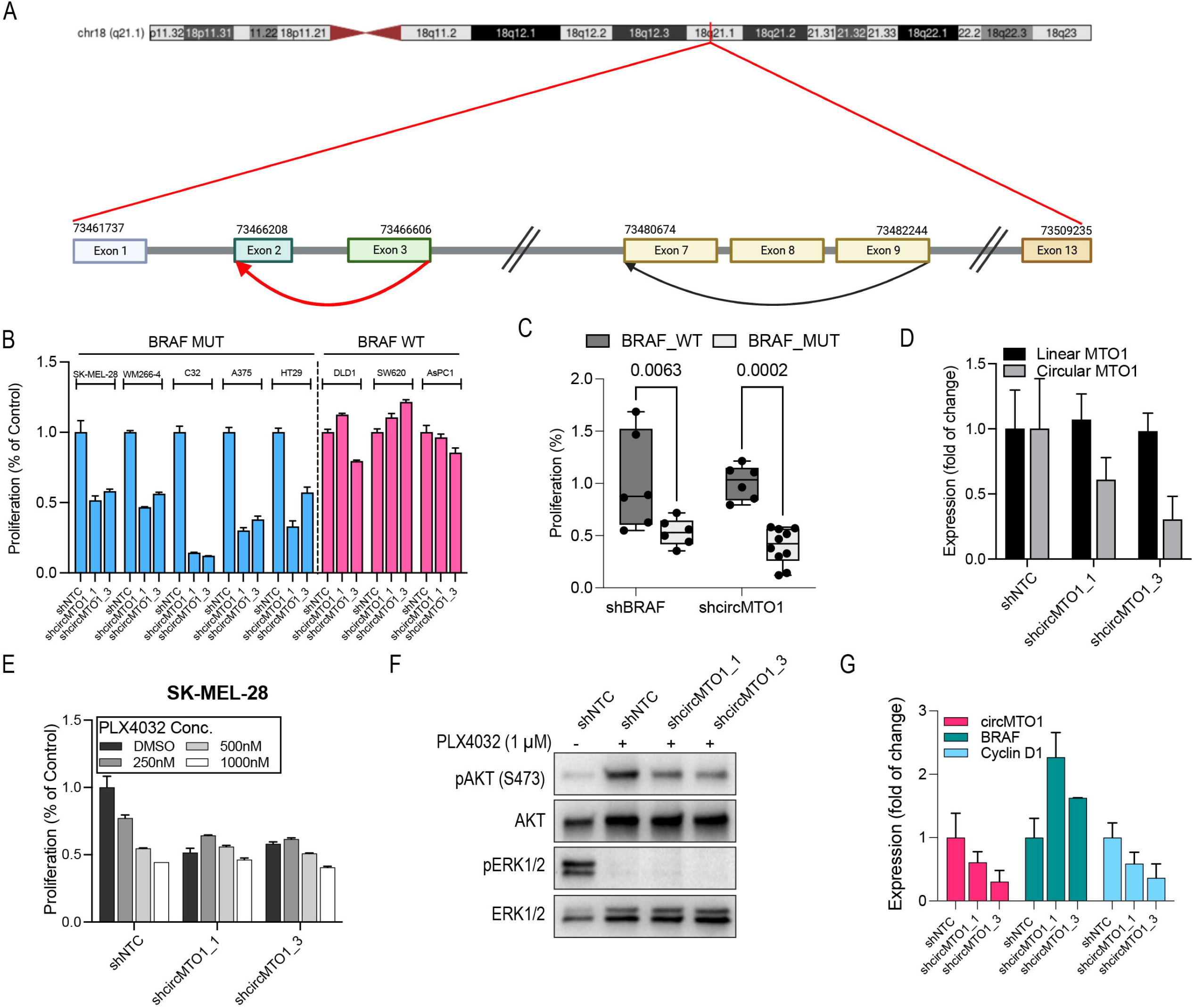
*circMTO1* is essential in BRAF mutated cancer cells. (A) Genome structure and circRNAs produced from *MTO1* gene. (B) Proliferation of *BRAF* mutant and WT cancer cell lines 7 DPI with shRNAs targeting *circMTO1*. (C) One-way Anova analysis of proliferation changes induced by *circMTO1* targeting shRNAs in *BRAF* mutant or WT cells. (D) Expression of circular and linear *circMTO1* in SK-MEL-28 cells 3 DPI with *circMTO1* targeting shRNAs. (E) Proliferation of SK-MEL-28 (*BRAF* mutated) following treatment with a BRAF inhibitor (PLX4032) and *circMTO1* targeting shRNAs showing no additive proliferation effect. (F) Levels of pERK and pAKT following treatment of SK-MEL-28 cells with PLX4032 and *circMTO1* targeting shRNAs. (G) mRNA levels of *circMTO1 BRAF* and *CCND1* mRNA following infection with *circMTO1* targeting shRNAs.

### circMTO1 regulates phosphorylation of ERK or AKT depending on PTEN status

To gain insight into the mechanisms of how *circMTO1* regulates the MAPK pathway, we looked at changes in the levels of pERK and pAKT, two key components of the MAPK pathway following shRNA mediated knockdown of *circMTO1*. We found that in *BRAF* WT cells knockdown of *circMTO1* upregulated pERK levels suggesting a feedback loop (Fig. 6A). Consistent with our observation that *circMTO1* is a regulator of MAPK signalling we found that in A375 and in Malme-3M, two *BRAF* mutated melanoma cell lines with high constitutive pERK, suppression of *circMTO1* inhibited pERK (Fig. 6A). Surprisingly, in three other *BRAF* mutated melanoma cell lines, (SK-MEL-28, C32 and WM266-4) suppression of *circMTO1* did not affect pERK levels but instead inhibited the levels of pAKT (Fig. 6A). This difference in the pathways that are regulated by *circMTO1* could be attributed to the levels of the tumour suppressor *PTEN* (Fig. 6A,B). In *BRAF* mutated cells with high PTEN levels and low pAKT (A375 and Malme-3M), depletion of *circMTO1* reduced pERK levels, while in *BRAF* mutated cells with low PTEN levels and high pAKT (SK-MEL-28, C32 and WM266-4), depletion of *circMTO1* reduced pAKT levels (Fig. 6A,B).

**Figure 6:**
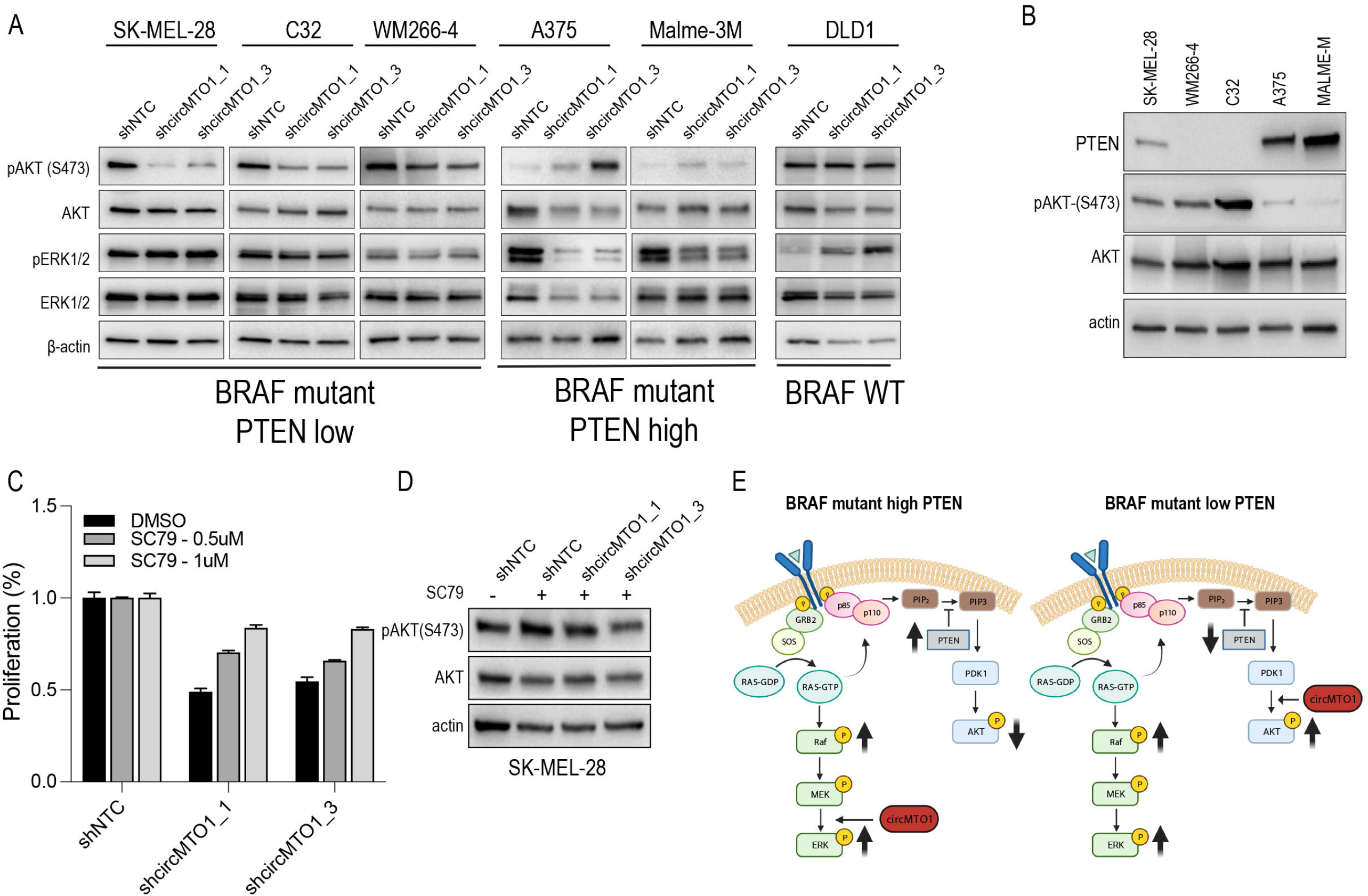
circMTO1 knockdown mediate suppression of pERK or pAKT in a PTEN dependent manner. (A) Levels of pAKT and pERK following knockdown of circMTO1 in a panel of BRAF mutated cancer cell lines. (B) Levels of PTEN and pAKT in the different BRAF mutated cell lines. (C) Rescue of proliferation in SK-MEL-28 following treatment with an AKT activator (SC79) and shRNAs targeting *circMTO1*. (D) pAKT protein levels following treatment with SC79 in SK-MEL-28 cells. (E) Suggested model of how *circMTO1* regulates proliferation in *BRAF* mutated cancers.

To validate these observations we used SC79, a molecule that binds to the PH domain of AKT and promotes its phosphorylation^42^. Following introduction of *circMTO1* targeting shRNAs in SK-MEL-28, cells were treated with DMSO or SC79 and proliferation was measured using a crystal violet staining assay. We observed a partial recue of proliferation arrest induced by *circMTO1* targeting shRNAs in *BRAF* mutated *PTEN*-low cells (Fig. 6C,D and Supplementary Fig. 6A,B), suggesting that circMTO1 regulation of pAKT activity is an important function of *circMTO1* in these cells. Our results demonstrate that *circMTO1* plays a key role in regulating the MAPK signalling pathway in a context specific manner that is dependent on *PTEN* status (Fig. 6E).

## Discussion

circRNAs are found throughout metazoans and are expressed in all cell types and organs. The extent to which circRNAs have functional roles in regulation of biological processes remains an open question, although circumstantial evidence including gene expression profiles and limited functional screens suggest that at least some circRNAs affect cell proliferation. Using capture arrays and RNASeq in >2,000 samples recent studies found that >100,000 circRNAs could be detected in human cells^2^. Most of these circRNAs are expressed at very low levels (in some cases only 1-2 reads in 1 sample), questioning if these low expressed circRNAs could be functional. Although circRNAs are very stable and are not degraded by the cellular RNA degradation machinery^43, 44^, a minimal number of circRNAs is likely to be required for mediating a function. In the current study we used a loss of function screen to evaluate biological processes and signalling pathways regulated by circRNAs. For these experiments we chose a minimal non-stringent expression cut-off (≥30 reads, ≥10 samples) for assessing functional circRNAs. Based in this criterion, we found 3,354 circRNAs that were used in functional screens.

Loss of function screens using gene targeting libraries have been highly successful in identifying genes associated with signalling networks^18^. Here, we applied a similar approach to identify circRNAs that function in a context specific or non-specific manner. The majority of circRNAs (70%) did not show any evidence of a phenotype following knockdown. This could be since most circRNAs are not functional or that the circRNAs have a role in processes that we did not measure or in a context that we did not include. Future screens that using more cell lines and phenotypes will be key for understanding the extent of functional circRNAs.

We found a subset of core essential circRNAs that are likely important for regulating core essential processes such as cell cycle, replication and apoptosis. Only a small number of circRNAs (176, 5% of circRNAs we tested) scored as core essential circRNAs and 20 (0.5%) validated in our secondary screen. This is consistent with the fact that only a small number of circRNAs are expressed at high levels across cell types^2^. We validated two of these circRNAs (*circRERE* and *circHUWE1*). Although both circRNAs validated as core essential and had a dramatic effect on cell proliferation, our data suggests that they have a different mechanism of action. Only knockdown of *circRERE* had an effect on proliferation and knockdown of linear *RERE* by CRISPRs or shRNAs had no effect on cell proliferation, suggesting they have distinct functions. Conversely, for *circHUWE1* we found that although knockdown of the circRNA did not affect the levels of linear *HUWE1*, suppression of *circHUWE1* or linear *HUWE1* had the same effect on cell proliferation. These observations suggest that in some cases circRNA may regulate a similar process as the linear mRNA. Although further characterisation of the mechanism of action of *circHUWE1* are needed these results suggest that at least in some cases the circular and linear RNAs regulate a similar cellular process.

The use of multiple cancer cell lines with different pathway dependencies allowed us to identify pathway specific circRNAs. In this study we investigated the mechanisms of circRNAs associated with MAPK and WNT pathways. This first proof of principle use of an “Achilles like” screen to identify pathway specific circRNAs demonstrates the utility of this approach. Additional studies are needed to identify other pathways that are likely to require other circRNAs. We also find some circRNAs that are lineage specific. This includes brain, colon, pancreas and skin specific circRNAs. Our validation studies show that these circRNAs are direct regulators of these pathways and suggest that *circMTO1* and *circSMAD2* may be used as targets for drug development and for understanding the molecular mechanisms that regulate the MAPK and WNT pathways in healthy or disease tissues.

Recent studies have suggested the use of CRISPR based approaches such as Cas13 to study circRNA function^45, 46^. Cas13 is a class of CRISPR enzymes that cleave RNA and could be repurposed for functional studies of non-coding RNAs. We have found three major issues that limit the applicability of CRISPR-Cas13 for functional studies of circRNAs. First, Cas13 ability to cut RNA is time sensitive. After a several days in culture Cas13 becomes inactive and thus it is difficult to observe long term phenotypes like proliferation. Second, RNAi usually shows a 70-90% reduction in circRNA expression but Cas13 usually shows much decreased levels of circRNA suppression (40-60%) making it difficult to identify phenotypes. Third, collateral damage activity^47^ induces a large number of off target effects and it is difficult to determine if an observed phenotype is a result of the on target or off target Cas13 effect. Although, the recent development of Cas7-11 with no observed collateral RNA cleavage may be used in future studies^48^, the current low cutting efficiency of the Cas7-11 system will need to be improved to enable functional readouts.

In summary, our study provides for the first time a comprehensive functional annotation of circRNAs. We provide a resource that will enable further detailed studies of how functional circRNAs mediate cellular processes. In this study we focused on circRNAs that regulate the WNT and MAPK pathways. Further studies will enable to uncover other circRNAs that are likely to regulate many other cellular functions.

## Methods

### Cell lines

Panc10.05 was gifted from Professor Roger Daly, Monash University, A2058 was gifted by Professor Grant McArthur, Peter MacCallum Cancer Centre, U87 was gifted from Dr. Paul Daniel, Hudson Institute of Medical Research, C32, WM266-4, SK-MEL-28 were gifted from Professor Mark Shackleton, Alfred Health. All other cell lines used in this study were from ATCC. HT29, DLD1, SW620, RKO, MIAPACA2 HPAC, A2058, Hs683, SF767, U87 were cultured in DMEM media containing 10% FBS, 1% Penicillin and Streptomycin and 1% Glutamine. COLO205, LS411N, ASPC1, A375 and LN229 were cultured in RPMI containing 10% FBS, 1% Penicillin and Streptomycin and 1% Glutamine.C32, WM266-4, SK-MEL-28 were maintained in DMEM media containing 10% FBS, 1% Penicillin and Streptomycin, 1% Glutamine and 1% NEAA. Malme-3M cells were cultured in IMDM media containing 20% FBS, 1% Penicillin and Streptomycin and 1% Glutamine. L3.3 was cultured in DMEM media supplemented with 10% FBS, 1% Penicillin and Streptomycin, 1% Glutamine, 1% NEAA and 1mM Sodium pyruvate. Panc10.05 was maintained in RPMI media containing 15% FBS, 1% Penicillin and Streptomycin, 1% Glutamine, 1.5 g/L Sodium bicarbonate, 4.5g/L glucose, 10mM HEPES, 1mM Sodium Pyruvate and 10 units/mL Insulin. IGR-39 was propagated in DMEM media supplemented with 20% FBS, 1% Penicillin and Streptomycin and 1% Glutamine. All cell lines were maintained in a humidified incubator at 37°C with 5% CO2.

### Generation of circRNA targeting shRNA library

shRNA sequences in pooled libraries are available in Supplementary Table 2 and 7. For each circRNA, we designed 5 shRNAs. shRNA pooled libraries were made as previously described^17, 49^. Briefly, oligo pools were purchased from custom arrays (WA, USA) containing shRNAs and flanking PCR handles with AgeI and EcoRI cutting sites. The final sequence of the oligo pool is: AACTTGAAAGTATTTCGATTTCTTGGCTTTATATATCTTGTGGAAAGGACGAAAC ACCGG(21nt SENSE)CTCGAG(21nt Anti SENSE)TTTTTGAATTCTCGACCTCGAGACAAATGGCAGTATTCATCCACAATTTTAAAAGAAAAGGGGG.

shRNA sequences were amplified from this pool with Fwd: 5’-AACTTGAAAGTATTTCGATTTCTTGGCTTTATATATCTTGTGGAAAGGACGAAAC ACCGG -3’, Rev: 5’-CCCCCTTTTCTTTTAAAATTGTGGATGAATACTGCCATTTGTCTCGAGGTCGAGAATTC – 3’ primers. The PCR product was cloned via Gibson assembly into AgeI and EcoRI digested pLKO.1 vector (Addgene#8453). Ligated libraries were electroporated into DH5a electrocompetent cells (Invitrogen), plasmid DNA was extracted using Qiagen Maxi Prep. For each library preparation, a 1000X representation was ensured.

### Loss of function screens

Cells were infected at an MOI of 0.3 at a multiplicity of 1,000 cells/shRNA. 24h post-infection, infected cells were selected using puromycin (2 μg/ml). To ensure shRNA expression, cells were maintained with puromycin throughout the screen. Twenty-one days post-infection, genomic DNA was extracted using NucleoSpin Blood XL (Clontech) and the shRNA sequence was amplified as previously described^50^. shRNA abundance was quantified using HiSeq Illumina sequencing. Deconvolution of shRNA read counts was done using the poolQ algorithm with default settings (https://portals.broadinstitute.org/gpp/public/software/poolq).

### RNA isolation and quantitative real-time PCR

Total RNA was isolated using TRIzol (Sigma Aldrich) from cell lines, according to the manufacturer’s instructions. 1μg of total RNA was reverse transcribed to cDNA using the Maxima Reverse Transcriptase (Thermo Fisher) in 20 μl reaction with random hexamers and dNTP. Expression levels of genes was quantified using qRT-PCR and the QuantStudio™5 Real-Time PCR system (Thermo Fisher). qRT-PCR was performed using the indicated qPCR primers (Supplementary Table 11). *GAPDH* was used for normalisation. The qRT-PCR conditions were denaturation at 95°C for 5 min followed by 40 cycles of amplification at 95°C for 15s and 60°C for 20s. The comparative cycle threshold (delta Ct) method was used to analyse the gene expression levels.

### Western blot analysis

Cells were washed in PBS and resuspended in RIPA buffer (CST#9806) containing proteinase inhibitors (Roche) and quantified using the Pierce BCA Protein Assay Kit (Thermo Fisher). Protein lysates diluted in 4 X Laemmli Sample Buffer (Bio-Rad#161-0747) were loaded onto Bio-Rad 4-20% precast gels. Following electrophoresis, proteins were transferred to a pre-activated PVDF membrane using the Trans-Blot®Turbo™ Transfer System and visualized using ECL (Bio-Rad Chemidoc). Antibodies used in this study are: GAPDH (Sana Cruz#SC32233), phospho-Akt (Ser473) (Cell Signalling#4060), Akt (Cell Signalling#4691), phospho-ERK1/2 (p44/p42) (Cell Signalling#4370), ERK1/2 (Cell Signalling#4695), β-actin (Sana Cruz #SC47778), PTEN (Cell Signalling#14672), CTNNB1 (Cell Signalling#8480).

### Crystal violet staining proliferation

Following shRNA infection and selection, cells were allowed to propagate for the indicated time. The media were removed, and cells were washed twice in PBS. 10% of formalin in PBS was added and incubated for 20 min at room temperature. Formalin was removed, and 0.5% (w/v) of crystal violet (Sigma#C0775-25G) was added and incubated for 20 min at room temperature. Crystal violet was removed, and plates were thoroughly washed with PBS. For quantification, 10% of acetic acid was added to each well and incubated at room temperature for 30 min. The extracted solution was added to a 96-well plate and quantified by measuring the OD at 590nm.

### Isolation and establishment of colon organoids (colonoids)

Mouse wildtype colon organoids (colonoids) were dissected following euthanasia and flushed multiple times with cold PBS. Adipose tissue was dissected off and mucous scraped away. The tissue was then sectioned into small pieces and washed further by multiple inversions in a tube until unattached epithelial fragments are separated. Intestinal tissue was then incubated in 0.5M EDTA on a spinning rotor machine at 10rpm at 4°C for 30 min. Samples were washed in PBS and vigorously pipetted the tissue to concentrate the crypts in the supernatant. Cells Crypt supernatant was centrifuged for 50 min (1,500 rpm) at 4°C. Cell pellet was resuspended in PBS and passed through a 70μM cell strainer and centrifuged for 5 min (1,500 rpm) at 4°C. Cell pellet were resuspended in Matrigel (∼400 crypts per 30μl of Matrigel) and seeded 30μl on a pre-warmed (37°C) 24-well plate. Colonoids were incubated and maintained in media containing: DMEM/F12 (Gibco#11320033), Glutamax 1% (Gibco#35050061), HEPES 1% (Gibco#15630130), Penicillin Streptomycin (1%) (Gibco#15140122), Fungizone (1%) (Gibco#15290018), B27 (1%), WNT3a conditioned media (50%), R-spondin conditioned media (10%), Noggin conditioned media (2%), Recombinant mouse EGF (1:1,000) (Peprotech#315-09). RKI-1447 (1:1,000) (In vitro technologies#1254) and CHIR99021 (1:3,000) (BioScientifica#04-0004) were added to the media for 24h during following passaging and seeding into Matrigel (BD#356231).

### Lentiviral transduction of colonoids

Colonoids were transduced with concentrated lentivirus as described by Maru et al.^37^. Specifically, 48h after seeding colonoids in media containing RKI-1447 (1:1,000) and CHIR99021 (1:3,000) hyperproliferative cystic crypts were liberated from Matrigel and resuspended in 250μl of 10X concentrated lentiviral organoid media with 8μg/ml polybrene. The colonoids in lentiviral organoid media and polybrene were then spun down on a bench top centrifuge at a room temperature for 1h (1,000rpm). Each colonoid infection condition was incubated in the organoid lentiviral media in a 37°C tissue culture incubator for 4h. Following the previously described infection procedure cells were spun down (1,500rpm) at 4°C to remove lentiviral media and the cell pellet was resuspended in ice cold Matrigel before seeding in a 48-well plate. Transduced colonoids were then maintained in media containing RKI-1447 (1:1,000) and CHIR99021 (1:3,000) for 48 hours, before being passaged onto a new plate. Colonoid media lacking RKI-1447 and CHIR99021 and containing 2μg/ml puromycin was added for selection of uninfected cells. Images were taken each day on an EVOS M5000 microscope to monitor colonoid growth.

### Lentiviral production

HEK293T cells were seeded in a 10-cm plate (6.5×10^6^ million cells per plate). 24h post seeding, cells were transfected using Lipofectamine™ 3000 reagent according to the manufacturer’s protocol. Transfection mix for each shRNA contained 8.75μg of shRNA targeting construct in pLKO.1 (Addgene#8453), 8.75μg psPAX2 (Addgene#12260) and 0.875μg of pCMV-VSV-G (Addgene#8454). Lentiviral media was collected at 24h and 48h timepoints and combined. Virus was concentrated using Amcion Ultra-15 30k centrifugal filter (Merck#C7715) and centrifugation for 30 min at 3,900rpm.

### WNT reporter activity assays

DLD1 cells were transduced with 7TFC (Addgene#24307) lentivirus. 48h post-transduction mCherry positive cells were sorted using a FACS Aria Fusion cell sorting machine. After 48h mCherry positive (7TFC transduced) cells were transduced with shRNA-targeting lentivirus and selected cells for 48h with 2 μg/ml puromycin. Selected cells were seeded at 5,000 cells/well in a 96-well plate, and spun cells down for 5 minutes (1,500rpm). 6h later, media was aspirated, and 15μl of lysis buffer was added (1mM HEPES, 5mM DTT, 2mM MgCl_2_, 2% v/v Triton X100). Plates were incubated for 10 min at room temperature and 100μl LAR Buffer (25mM Glycylglycine, 15mM K_x_PO_4_ (pH=8), 4mM EGTA, 2mM ATP, 1mM DTT, 15mM MgSO4, 0.1mM CoA and 75μM D-luciferin). Luminescence was measured on a BMG Pherastar microplate reader and normalised to mCherry.

### Cell viability assay

RKO cells were seeded at 300,000 cells/well on a 6-well plate and transduced with either shNTC or shcircRERE lentivirus. Following 24h incubation lentiviral media was removed and replaced with complete media containing 2μg/ml of puromycin. Cells were selected in puromycin media until non-transduced cells were all dead (∼24h). After selection, transduced cells were plated on a 12-well plate at 50,000 cells/well with or without different drug treatments. Following a 24h incubation media was replenished with fresh media containing the same drug concentration or DMSO. After 48h of drug treatment, total well contents (both floating and attached cells) for each drug treatment was collected and stained with trypan blue dye (1:1 ratio cells:trypan blue dye). Live and dead cell counts were measured using an Invitrogen Countess 3 Automated Cell Counter.

### Cell viability confocal microscopy assay

RKO cells were seeded at 300,000 cells/well on a 6-well plate and transduced with either shNTC or shcircRERE lentivirus. Following 24h incubation lentivirus, media was removed and replaced with complete media containing 2μg/ml of puromycin. Cells were selected in puromycin media until non-transduced cells were all dead (∼24h). After selection, 5,000 cells/well were plated onto an LabTek 8-well chamber slide (ThermoFisher#NUN155411). Media in each chamber contained puromycin with DMSO plus/minus different drug treatments for shcircRERE infected cells. Media was replenished with drug treatments at the same concentration every 24h for 48h. After 48h of drug treatment cells were stained cells with Propidium iodide (PI) (2ug/ml) and Fluo4AM (1uM) in complete DMEM media for 30 min in a 37°C incubator. After drug treatment, media was removed and replaced with complete DMEM. Cells were visualised using a laser scanning confocal microscope (LSM780, Zeiss) with water UV-VIS-IR Apochromat 40X 1.20W Korr FCS M27 objective, Immersion Oil W 2010 (Zeiss, 444969-0000-000) and avalanche photodiode light detectors (APDs) of the Confocor 3 module, fitted with an incubation chamber for controlled conditions at 37°C and 5% CO_2_. GaAsP detectors were used with the 488 and 561 laser lines combined in the same track with appropriate filter settings depending on the fluorophores. 3D images of cells were acquired using the following parameters: image size 512 × 512, pinhole 2uM, pixel dwell time 6.3 μm/s, at ∼0.4μm section intervals.

### Image and statistical analysis

Image analyses were performed using Imaris 7.4.2 and 9.5.1 software (Bitplane AG) and ImageJ/Fiji software. Mean cell fluorescence intensities of individual cells was measured on 3D surfaces using the Fiji measure function. Statistical analyses were performed using GraphPad Prism software.

## Acknowledgments

This work was supported by an Australian Research Council (ARC) grant to J.R., G.G. and P.T. (grant number: DP210101755). J.R is supported by a Victoria Cancer Agency fellowship (grant number: MCRF20035).

We thank the Functional Genomics Platform, the Bioinformatics Platform and Micromon genomics platform at Monash University for help with loss of function screens and data analysis. We thank Dr. Thierry Jarde and Prof. Helen Abud from the Monash organoid platform for help with organoid experiments. We thank Dr. Michael Lazarou from the Walter and Eliza Hall Institute for helpful discussions and suggestions.

## Supplementary Figure Legends

**Supplementary Figure 1: Primary screen identifies context specific and non-specific circRNAs**. (A) Overlap between circRNAs profiled in this study and circRNAs that were profiled in previous prostate specific circRNA screens^9^. (B) Distribution of proliferation changes following shRNA mediated knockdown of core essential genes. (C) Two class comparisons identifying pathway and lineage specific circRNAs.

**Supplementary Figure 2: Secondary screen validates context specific and non-specific circRNAs**. Distribution of proliferation changes following shRNA mediated knockdown of core essential genes.

**Supplementary Figure 3: Identification of core essential circRNAs**. (A) Genome structure and circRNAs produced from *HUWE1* gene. (B) Proliferation changes observed in primary screen following suppression of *RPS8* (cell essential gene, positive control) or two *HUWE1* derived circRNAs. (C) Validation of HUWE1_chrX_53614533_53615835 in secondary proliferation screens. (D) Expression of circular or linear *HUWE1* following introduction of *circHUWE1* targeting shRNAs in DLD1 cells. (E) Crystal violet staining in a panel of cell lines following knockdown of *circHUWE1*. (F) Quantification of proliferation changes from (E). (G) Proliferation changes from Depmap^51^ following CRISPR mediated suppression of *HUWE1* gene. (H) Proliferation changes from Depmap^27^ following shRNA mediated suppression of *HUWE1* gene. (I) Crystal violet staining in a panel of cell lines following knockdown of *circRERE*. (J) Cell viability was assessed using trypan blue staining in RKO cells following transduction with *shcircRERE_1* or *shNTC* or *shcircRERE_1* containing cells were treated with either 4μM of Ferrostatin-1 or 10μM of Necrostatin or 20μM of QVD-OPH. Cell viability was measured 48h after drug treatment.

**Supplementary Figure 4: Identification of circRNAs that regulate the WNT signalling pathway**. (A) Crystal violet images following introduction of shRNAs targeting *circSMAD2* in WNT active or inactive cell lines. (B) Crystal violet images following introduction of shRNAs targeting *circZNF652* in WNT active or inactive cell lines. (C) Quantification of crystal violet from (B). (D) Crystal violet images following introduction of shRNAs targeting *circPKHD1* in WNT active or inactive cell lines. (E) Quantification of crystal violet from (D). (F) Expression of linear or circular *ZNF652* following transduction of *circZNF652* shRNAs in DLD1 cells. (G) Expression of linear or circular *PKHD1* following transduction of *circPKHD1* shRNAs in DLD1 cells. (H) Crystal violet images of rescue of *circSMAD2* induced proliferation inhibition following over expression of *CTNNB1* in a WNT active (DLD1) or WNT inactive (RKO) cell line.

**Supplementary Figure 5: Validation of circRNAs that are essential for KRAS mutant cell lines**. (A) Crystal violet images of KRAS WT or mutant cells infected with *circRNF170* targeting shRNAs. (B) Quantification of proliferation changes in (A). (C) Expression of *circRNF140* in MIAPACA2 cells following shRNA transduction. (D) Crystal violet images of KRAS WT or mutant cells infected with *circNOL12* targeting shRNAs. (E) Quantification of proliferation changes in (D). (F) Expression of *circNOL12* in MIAPACA2 cells following shRNA transduction. (G) Crystal violet images of KRAS WT or mutant cells infected with *circNUDC* targeting shRNAs. (H) Quantification of proliferation changes in (G). (I) Expression of *circNUDC* in MIAPACA2 cells following shRNA transduction. (J) Comparison of proliferation changes induced by shRNAs targeting *circRNF140, circNOL12* or *circNUDC* in KRAS WT or mutant cells. One way anova analysis was used to calculate a pValue. (K) Levels of pERK and pAKT following suppression of *circRNF170* in KRAS WT and KRAS mutant cells. (L) Levels of pERK and pAKT following suppression of *circRNF170* in KRAS WT and KRAS mutant cells.

**Supplementary Figure 6: Identification of circMTO1 as a regulator of BRAF dependent cancers**. (A) Crystal violet images following introduction of shRNAs targeting *circMTO1* in *BRAF* dependent or independent cell lines. (B) Proliferation of C32 (*BRAF* mutated) cells following treatment with a BRAF inhibitor (PLX4032) and *circMTO1* targeting shRNAs showing no additive proliferation effect. (C) Proliferation of WM266-4 (*BRAF* mutated) cells following treatment with a BRAF inhibitor (PLX4032) and *circMTO1* targeting shRNAs showing no additive proliferation effect.

**Supplementary Figure 7: Knockdown of circMTO1 inhibits pERK or pAKT depending on the levels of PTEN**. (A) Partial rescue of proliferation in WM266-4 cells following treatment with an AKT activator (SC79) and shRNAs targeting *circMTO1*. (B) Rescue of circMTO1 inhibition in pAKT levels following treatment with SC79 in WM266-4 cells.

## Supplementary Tables

**Supplementary Table S1: Genomic features of circRNAs that are used in this study**. Genomic coordinates and read counts of circRNAs that were used in this study.

**Supplementary Table S2: Sequences of shRNAs used in pooled primary screens**. Target sequences of shRNAs targeting circRNAs or core essential genes that were used in primary screens.

**Supplementary Table S3: Read numbers from primary screen**. Read numbers in shRNA pool or in cells infected with circRNA targeting library 21 days post infection.

**Supplementary Table S4: Fold change calculations in primary circRNA screen**. Log2 Fold change in circRNA abundance was calculated by comparing shRNA abundance after 21 days in culture to the DNA pool and using the average of all 5 shRNAs targeting each circRNA.

**Supplementary Table S5: Correlation between gene dependency from Depmap and circRNA dependency**. For each circRNA Pearson was used to calculate correlation between the proliferation effect of the linear (CRISPR or shRNA from^27, 51^) and circRNA on proliferation.

**Supplementary Table S6: Genes that score in two class comparisons in primary screens**. For each two-class comparison, the difference between classes was determined by calculating the difference in viability between two classes.

**Supplementary Table S7: Sequences of shRNAs used in pooled secondary screens**. Target sequences of shRNAs targeting circRNAs or control genes that were used in secondary validation screens.

**Supplementary Table S8: Read numbers from secondary screen**. Read numbers in shRNA pool or in cells infected with circRNA targeting library 21 days post infection.

**Supplementary Table S9: Fold change calculations in secondary circRNA screen**. Log2 Fold change in circRNA abundance was calculated by comparing

shRNA abundance after 21 days in culture to the DNA pool and using the average of all 5 shRNAs targeting each circRNA.

**Supplementary Table S10: Genes that score in two class comparisons in secondary screens**. For each two-class comparison, the difference between classes was determined by calculating the difference in viability between two classes.

**Supplementary Table S11: Primers and shRNAs used in this study**.

